# YOLOv8 Enables Automated Dragonfly Species Classification Using Wing Images

**DOI:** 10.1101/2024.09.01.610694

**Authors:** Suyeon Kim

**Affiliations:** Greenfingers Jeju, Daejeong-eup Edu City, Seogwipo-si, Jeju Special Self-Governing Province, Korea

**Author notes:** Corresponding author: Suyeon Kim.

**Keywords:** Insect collection, Auto digitization, Auto classification, Computer vision, YOLOv8

## Abstract

This study investigates the digitization of insect collections and the application of deep learning models to improve this process. During a one-hour filming session, 141 images of dragonfly specimens from the Cornell University Insect Collection were captured and preprocessed using five distinct methods: (1) adding box annotations around the wings, (2) adding polygon annotations to outline the forewings and hindwings, (3) removing vein system images, (4) retaining only the wing outline images, and (5) grouping by automatically measured wing size and classifying species within those groups. By comparing YOLOv8 models trained on datasets with these different preprocessing methods, the study revealed three key findings: (1) datasets with bounding box annotations result in shorter preprocessing times and superior model performance compared to polygon annotations; (2) although models trained with polygon annotations may have lower accuracy, they provide more detailed information on wing length and phenotypic traits; and (3) the wing vein system, rather than the wing outline, is the critical factor in classification accuracy.

## 1. Introduction

In any insect-related study, one of the primary tasks is the collection of specimens, which requires careful preservation for future research (Kumar et al. 2022). However, as the use of these collections grows, insect collections worldwide are increasingly challenged in maintaining the original, undamaged condition of their specimens (Schauff 2001). Additionally, the analog format of these collections limits their accessibility to most researchers and potential users (Hudson et al. 2015). As a result, these collections often remain unused and obscure, known only to a small group of specialists associated with the institutions that house them (Hudson et al. 2015).

A globally emerging solution to these challenges is the digitization of collection datasets, combined with the application of computer vision and deep learning models to enhance time efficiency (Leusen and Heerlien, year; Høye et al., 2021). The digital format of insect collections not only allows for the permanent preservation of each specimen’s physical appearance, along with additional biological information, but also enables the quantification of variation in phenotypic traits and behavior, thereby preventing deterioration (Høye et al., 2021).

Constructing a quick and precise classification model is a crucial first step in digitizing visual and biological information from specimens in their pristine state and analyzing it for further research. This can be achieved through the application of deep learning artificial intelligence, which can automatically detect meaningful patterns in data (Høye et al., 2021). By digitizing taxonomic features and building a classifier model, researchers have a promising tool for accurately classifying and identifying plants and animals (Yang et al., 2015; Barré et al., 2017; Tuda and Luna-Maldonado, 2020).

A study on the InsectNet deep learning model, trained on 6 million images from iNaturalist, achieved over 96% accuracy in classifying 2,526 insect species using a self-supervised learning approach, highlighting its potential to reduce training time and costs (Chiranjeevi et al. 2024). Another study used web-crawled images to train a residual neural network on 94 dragonfly species, achieving 61.35% accuracy on test data (Balaji et al. 2022). However, these approaches require about 80 images per species, which tends to bias the model toward more commonly distributed species. There have been fewer attempts to classify specimens with only a limited number of samples, such as those housed in museums, compared to efforts with large-scale datasets.

Dragonflies have significant potential for classification based on their wings. Chitsaz et al. (2020) conducted a study examining the geometric parameters of photogrammetrically reconstructed wings from 75 Odonata species—66 from Epiprocta and 9 from Zygoptera. They found that wing shape and aero-structural characteristics, which influence key behaviors, are distinct for each species. The study revealed that features such as wingspan length, chord lengths, nodus location, and pterostigma size are independent across the Odonata class, highlighting the unique structural attributes of dragonfly wings.

If the wing structure of dragonflies can significantly enhance the accuracy of models trained on small datasets to a level comparable with large dataset models, future digitization efforts could greatly reduce the time required for data collection, preprocessing, and annotation. Additionally, archiving the biological details of dragonfly wings could provide valuable support for future ecological research. Therefore, this study aimed to identify an effective and economical approach to building an automated classification model for dragonflies without relying on large datasets.

## 2. Methods and Materials

### CAMERA DEVELOPMENT

**Figure 1.**
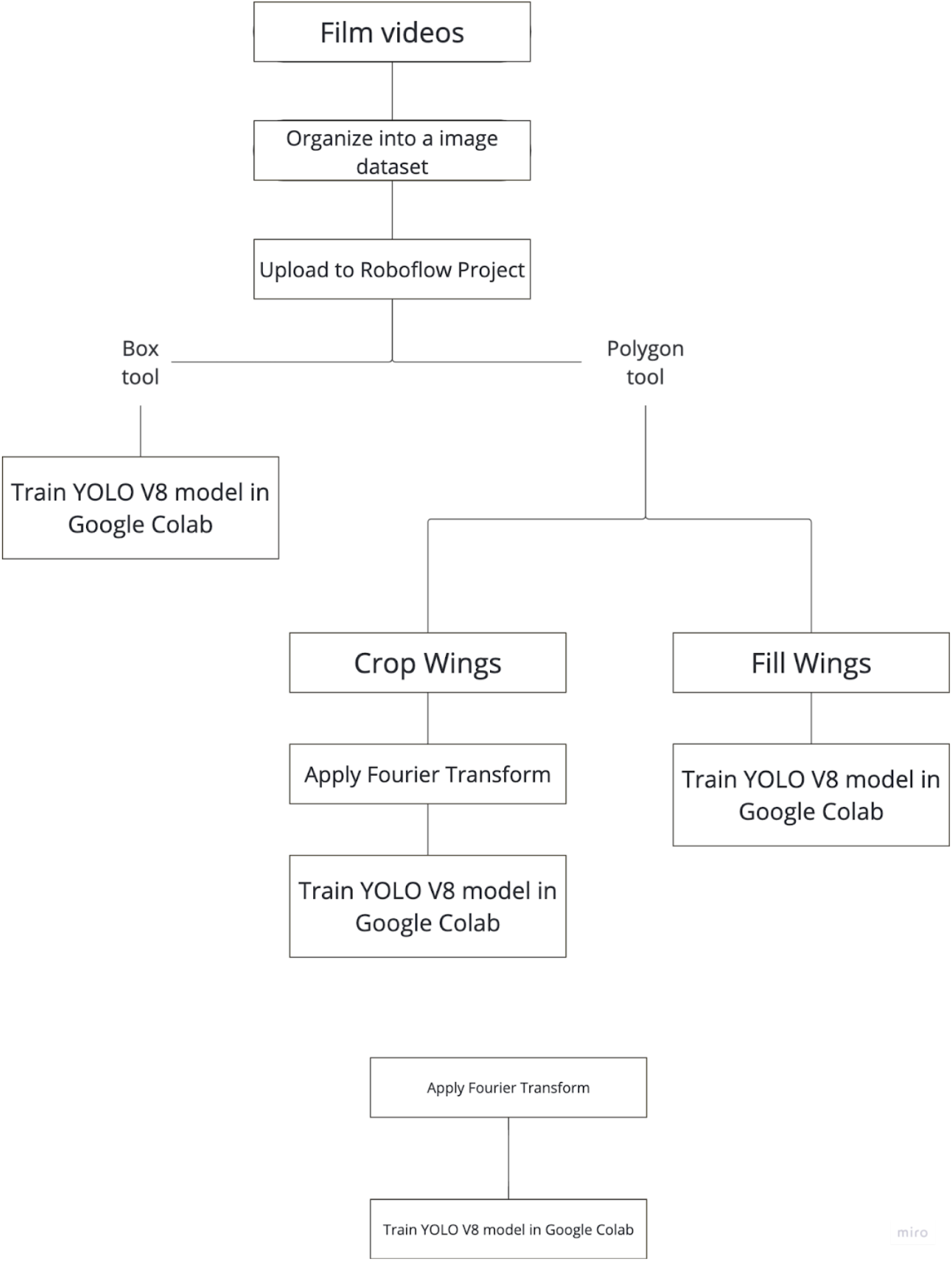
The image processing process. **Film videos:** The process begins with recording videos. **Organize into an image dataset:** The videos are then organized into an image dataset. **Upload to Roboflow Project:** The dataset is uploaded to a Roboflow project. **Box tool:** This method is used for simpler annotations, which involves creating a bounding box around the wings. The next step is to train a YOLO V8 model. **Polygon tool:** This method involves more detailed annotations using polygon shapes. **Crop Wings:** The wings are cropped out, and a Fourier Transform is applied. This data is then used to train a model. **Fill Wings:** The entire inner section of the wings is filled, leaving only the outline. This data is also used to train a model. **Additional Training:** The Fourier Transform is applied again, followed by further training of a model.

**Figure 2.**
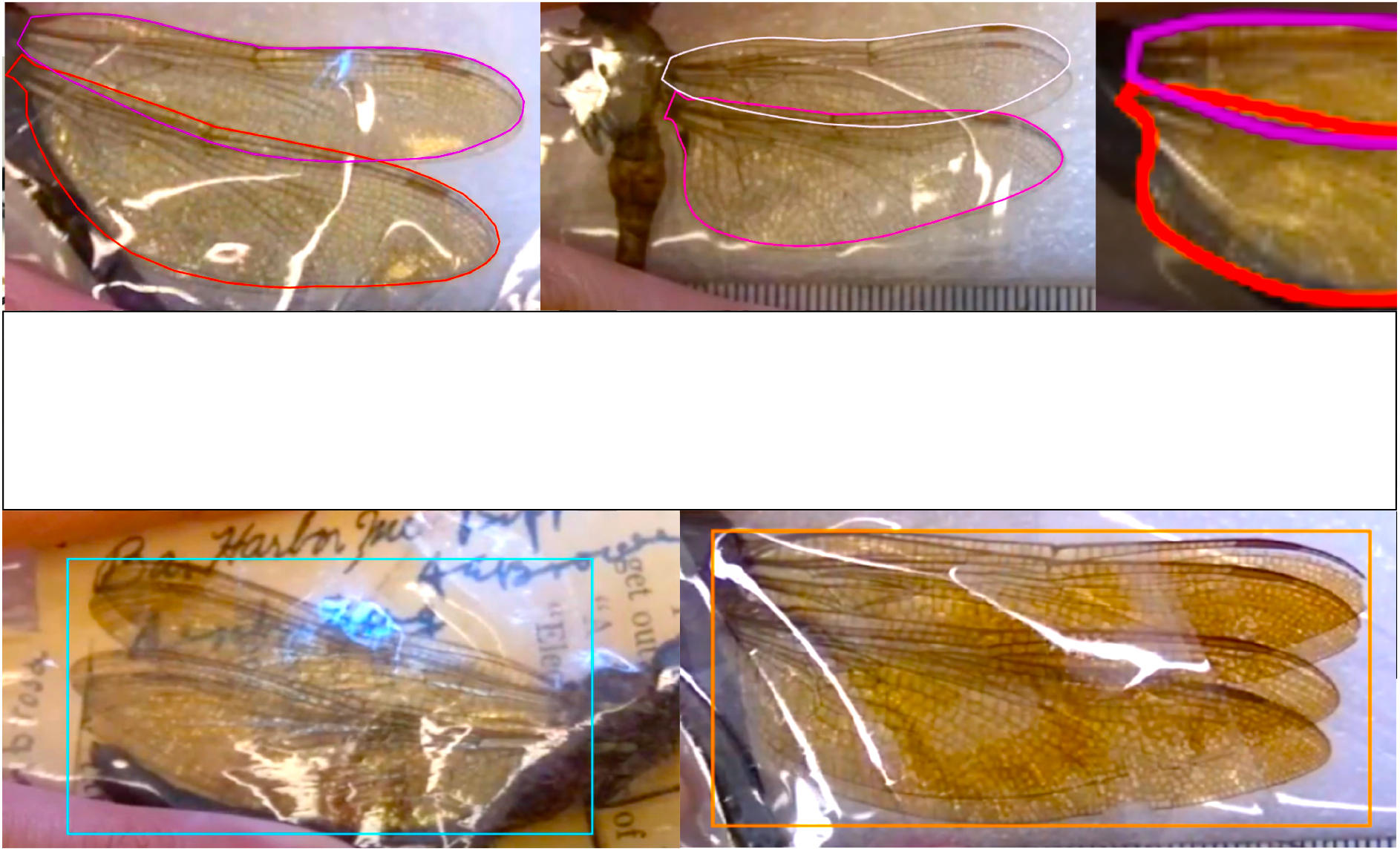
Annotation following the outline of the dragonfly wings (Above). Annotation with a rectangular box including the wings and background (Below).

This study utilized two units of Raspberry Pi camera setups, each comprising one HQ Camera (Arducam 8-50mm C-Mount Zoom Lens), HQ module (IMX477), Raspberry Pi 4 Model B, Romoss 30W 20000mAh power bank, and a tripod. A portable monitor was connected to the Raspberry Pi 4 Model B through a micro HDMI to Type B cable when needed.

### DATA COLLECTION

Dragonfly specimens were photographed at the Cornell University Insect Collection (CUIC). The species used in the study were selected based on the following criteria: at least 24 specimens per species, with the first six species meeting this criterion chosen from specimen boxes arranged alphabetically from A to Z. The specimens were stored in plastic envelopes with their wings folded, and the wing photographs were taken without removing the specimens from these envelopes. On average, it took 10 seconds to record the name, voucher, and photograph the wings of each specimen.

To acquire the images of the specimens, the HQ camera was positioned perpendicularly above the desk to capture the entire surface of the dragonfly wings. The HQ camera was activated through “libcamera” commands on the Raspberry Pi 4 terminal. While filming, the left wings were placed above the right wings, except for a few dragonflies whose left wings could not be separated from the paper labels. Specimens with heavily damaged wings were omitted. The videos were stored as h264 files on the Raspberry Pi 4 and later manually screen-captured to obtain images of the dragonfly wings as data images.

### ANNOTATING AND IMAGE PRE-PROCESSING

The captured images were uploaded to a Roboflow Object Detection Project. Roboflow provides two types of annotation tools: the polygon tool and the bounding box tool. To compare the performance of these two annotation methods, a copy of the entire dataset was manually annotated using the polygon tool. For the bounding box tool, another copy of the dataset was used, with only 100 images manually annotated to activate the auto-labeling function. The rest of the images were then manually approved or adjusted after being automatically labeled and annotated. The Grayscale and Auto-Orient options were applied to both copies of the datasets.

When using the polygon tool, classes were created in the form of “species-fore” or “species-back,” each indicating the forewing or backwing of a certain species, and specific rules were followed for outlining. If the wing was undamaged and too curvy to be perfectly represented by the polygon tool, it was preferable to exclude tiny parts of the wing rather than include background elements. However, if the wing was slightly damaged, it was better to allow some inclusion of the background to make a complete wing shape. Additionally, the outline had to include transparent parts only; dark copulas between the body and wing were excluded as much as possible.

For the bounding box method, both wings were enclosed in a single box without distinguishing between the forewing and backwing.

### DEEP IMAGE PRE-PROCESSING

**Figure 4.**
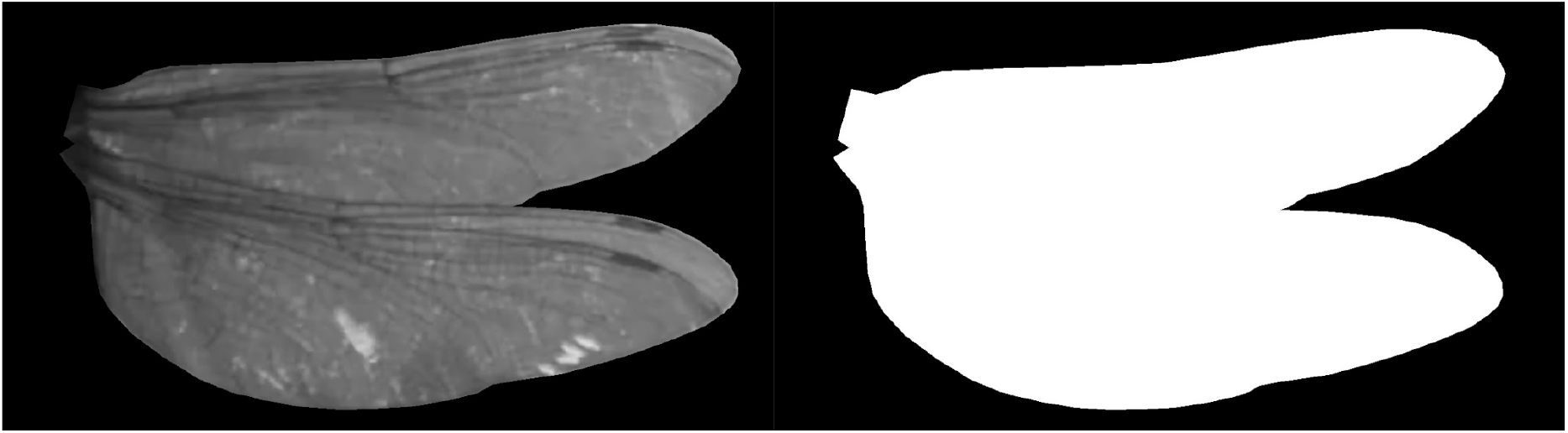
The first method removes the background, retaining the wing’s internal structure (veins; Left). The second method fills the wing’s interior with white, keeping only the outline (Light).

To compare whether the model finds the wing outline or the wing’s internal veins more useful for classification, two preprocessing methods were separately applied to two copies of the entire dataset. The first method involved removing all the background outside the wing outline while retaining the internal wing structure (i.e., the veins). The second method involved filling the entire inner section of wing with white, leaving only the outline visible in the image.

### MODEL DEVELOPMENT

Using the functions provided by the ultralytics library in google colab environment, multiple YOLO V8 model trained with differetly processed dragonfly wing images were developed (shared in Appendix). The YOLO model trained with images annotated by the polygon tool was expected to detect the wing outline, classify species, and distinguish between forewings and backwings. The objective of the model trained with the bounding box dataset was to detect the presence of wings and classify species. The train, validation, and test split ratios were 70:15:15, and accuracy was evaluated using precision, recall, mAP50, and mAP50-95.

### WING AREA AND LENGTH MEASURMENT

**Figure 5.**
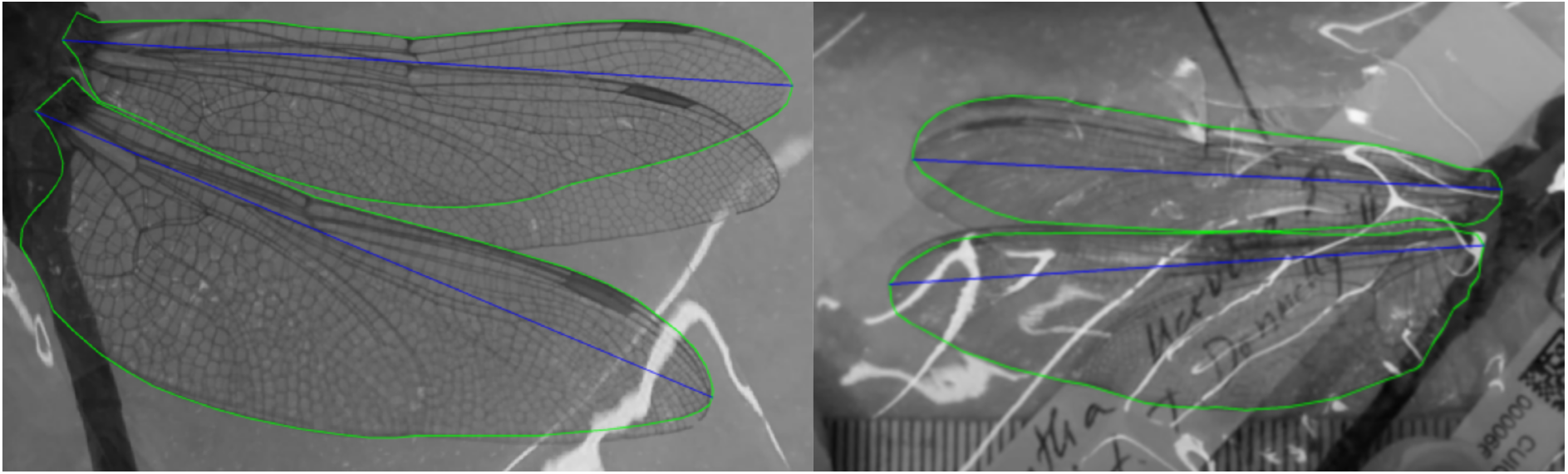
Examples of wing length (blue lines) measurment.

This study investigated whether adding information on wing area and length measured using YOLO could improve classification performance. To obtain accurate measurements, a ruler was photographed at a fixed camera height and angle to count the number of pixels per centimeter as a reference. The length of the wing was defined as the longest line drawn between two points on the wing outline, and the centimeter length was estimated from the pixel count of this line. The wing area was calculated by counting the number of pixels within the wing outline.

Next, this study developed a YOLOv8 model that first classifies dragonflies into three groups based on wing length. Subsequently, the model performs further classification within each group using the wing images.

## 3. Results and Discussion

### 3.1 Comparison of Annotation Methods: Bounding Box vs Wing Outlining

**Figure 6.**
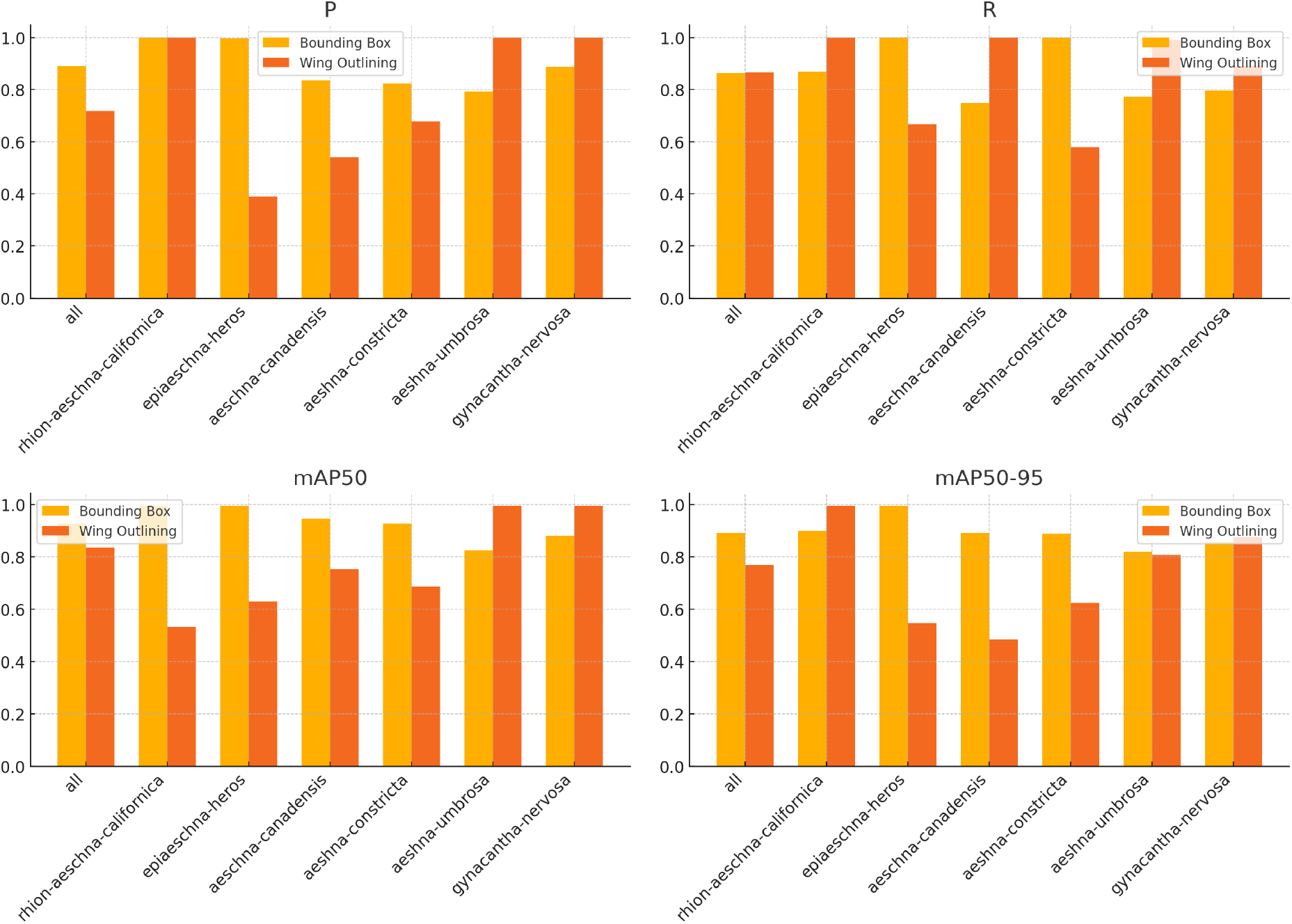
A comparision of species identification performance with bounding box vs. wing outlining annotation.

The bounding box dataset exhibited higher overall recall (0.865) and precision (0.89) compared to the polygon dataset. Bounding box results for all species showed reliable precision and recall (at least 75%). In contrast, the polygon dataset had species with precision as low as 30%.

For Rhion-aeschna-californica species, the bounding box model produced higher precision (1.0) and recall (0.869). For the polygon dataset, the recall was 0.793 for back wings and 1.0 for forewings, with precision values of 0.667 for back wings and 0.532 for forewings. For Epiaeschna-heros species, the bounding box model produced higher precision (0.998) and recall (1.0) compared to the polygon model, which had a precision of 0.542 for forewings and 0.492 for hindwings, with recall values of 1.0 and 1.0, respectively. For Aeschna-canadensis species, the polygon tool model produced higher recall (0.902 for forewings and 0.902 for hindwings) and similar precision (1.0 for forewings and hindwings) compared to the bounding box model, which had a precision of 0.836 and recall of 0.75. For Aeshna-constricta species, the bounding box model produced higher recall (1.0 for both) and precision (0.824 for bounding box compared to 1.0 for polygon forewings and hindwings). For Aeshna-umbrosa species, the bounding box model produced higher recall (0.774) and precision (0.793) compared to the polygon model, which had a precision of 0.542 for forewings and 0.309 for hindwings, with recall values of 0.667 and 0.6, respectively. For Gynacantha-nervosa species, both models produced similar results for recall (0.797 for bounding box and 0.66 for polygon) and precision (0.889 for bounding box and 0.713 for polygon forewings, 0.68 for hindwings).

It took approximately 10 minutes per sample to annotate the boundary of the wings, whereas annotating in the bounding box format took about 10 seconds per sample. In terms of efficiency, the box annotation was about 60 times more efficient.

**Table 2.**
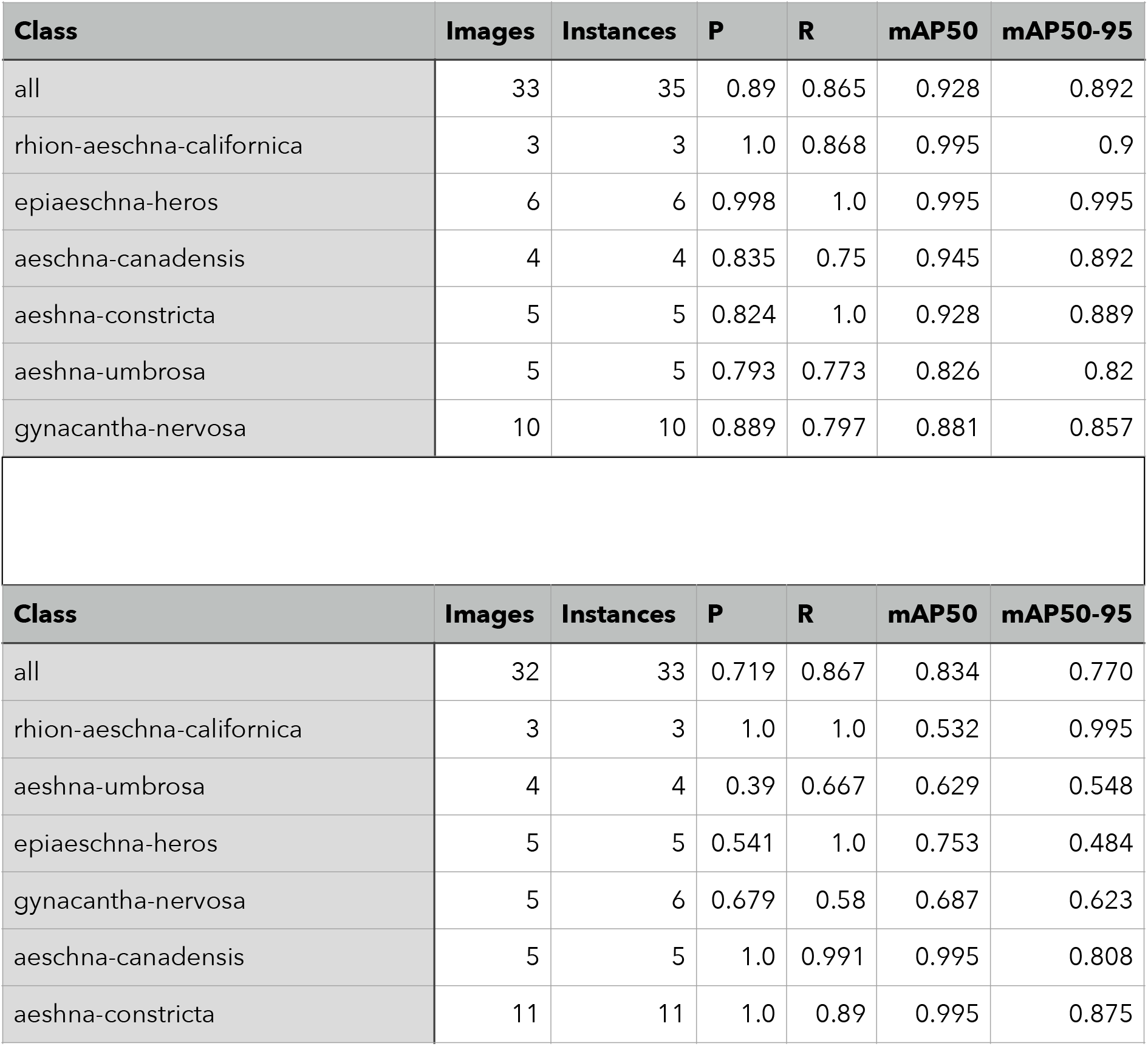
Species classification performance with bounding box annotation (Above) and wing outlining annotation (Below).

| Class | Images | Instances | P | R | mAP50 | mAP50-95 |
| --- | --- | --- | --- | --- | --- | --- |
| all | 33 | 35 | 0.89 | 0.865 | 0.928 | 0.892 |
| rhion-aeschna-californica | 3 | 3 | 1.0 | 0.868 | 0.995 | 0.9 |
| epiaeschna-heros | 6 | 6 | 0.998 | 1.0 | 0.995 | 0.995 |
| aeschna-canadensis | 4 | 4 | 0.835 | 0.75 | 0.945 | 0.892 |
| aeshna-constricta | 5 | 5 | 0.824 | 1.0 | 0.928 | 0.889 |
| aeshna-umbrosa | 5 | 5 | 0.793 | 0.773 | 0.826 | 0.82 |
| gynacantha-nervosa | 10 | 10 | 0.889 | 0.797 | 0.881 | 0.857 |

| Class | Images | Instances | P | R | mAP50 | mAP50-95 |
| --- | --- | --- | --- | --- | --- | --- |
| all | 32 | 33 | 0.719 | 0.867 | 0.834 | 0.770 |
| rhion-aeschna-californica | 3 | 3 | 1.0 | 1.0 | 0.532 | 0.995 |
| aeshna-umbrosa | 4 | 4 | 0.39 | 0.667 | 0.629 | 0.548 |
| epiaeschna-heros | 5 | 5 | 0.541 | 1.0 | 0.753 | 0.484 |
| gynacantha-nervosa | 5 | 6 | 0.679 | 0.58 | 0.687 | 0.623 |
| aeschna-canadensis | 5 | 5 | 1.0 | 0.991 | 0.995 | 0.808 |
| aeshna-constricta | 11 | 11 | 1.0 | 0.89 | 0.995 | 0.875 |

### 3.2 Comparison of Annotation Methods: Wing Outline vs Wing Veins

**Figure 7.**
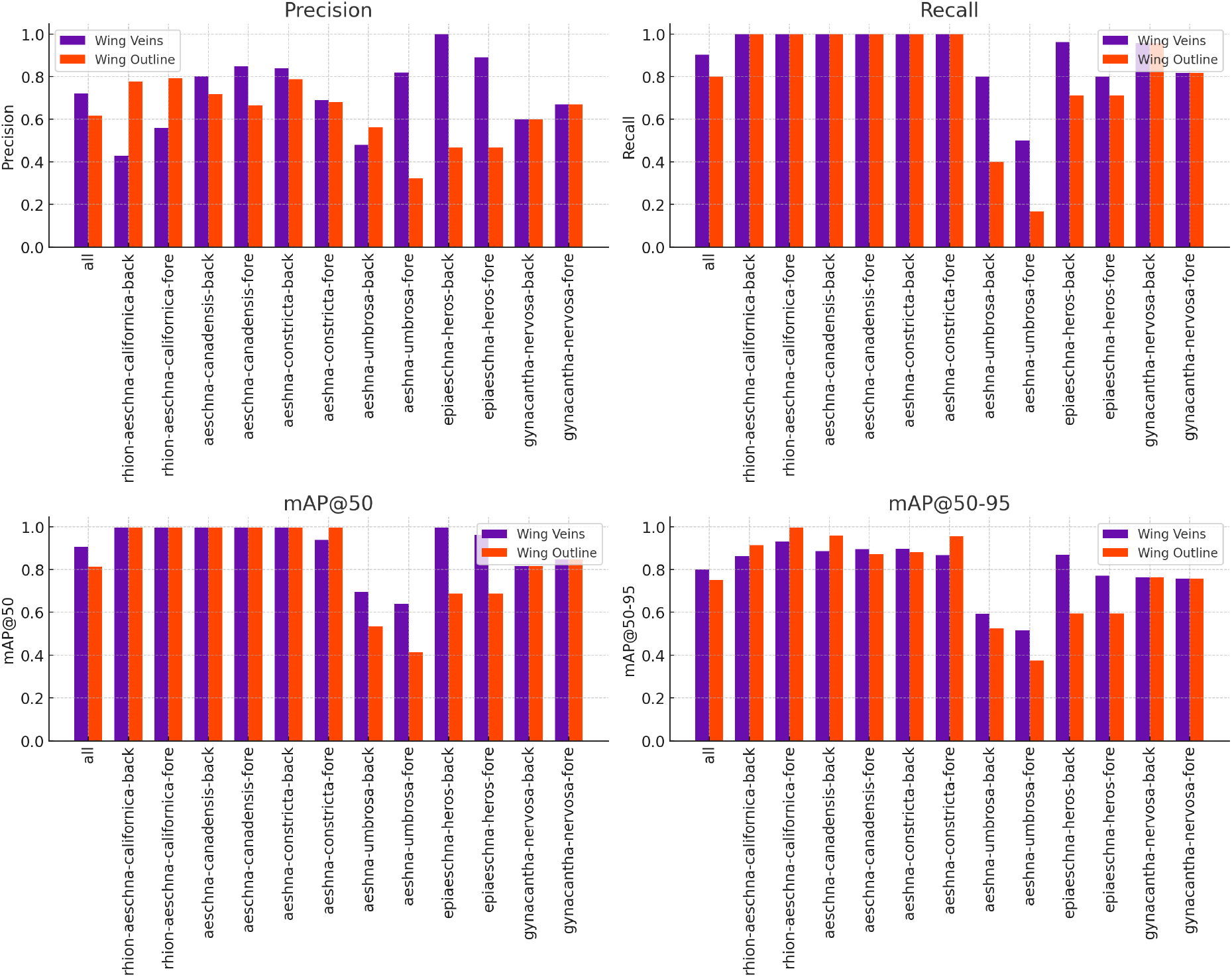
A comparision of species identification performance with wing vein information or wing outline information.

To compare whether the model finds the wing outline or the wing veins more useful for classification, two preprocessing methods were separately applied to two copies of the entire dataset.

In terms of Precision, wing-vein information shows higher precision in the overall class (“all”), as well as in classes like *aeschna-canadensis-back, aeschna-canadensis-fore, aeshna-constricta-back, aeshna-constricta-fore, aeshna-umbrosa-fore, epiaeschna-heros-back*, and *epiaeschna-heros-fore*. However, wing-outline information outperforms wing-vein information in classes such as *-rhion-aeschna-californica-back, -rhion-aeschna-californica-fore, aeshna-umbrosa-back*, and *gynacantha-nervosa-back*.

In terms of Recall, wing-vein information generally achieves higher or equal recall compared to wing-outline, particularly in the overall class (“all”) and in most specific classes like *aeshna-umbrosa-back, aeshna-umbrosa-fore, epiaeschna-heros-back*, and *epiaeschna-heros-fore*. The recall is equal in several classes (*-rhion-aeschna-californica-back, -rhion-aeschna-californica-fore, aeschna-canadensis-back, aeschna-canadensis-fore, aeshna-constricta-back, aeshna-constricta-fore*) with a perfect recall value of 1.0 in both treatments.

In terms of mAP50, wing-vein information generally outperforms wing-outline in mAP50 across most classes, including the overall class (“all”), and classes such as *aeshna-umbrosa-back, aeshna-umbrosa-fore, epiaeschna-heros-back, epiaeschna-heros-fore, gynacantha-nervosa-back*, and *gynacantha-nervosa-fore*. However, wing-outline information shows slightly better performance in the *-rhion-aeschna-californica-back, -rhion-aeschna-californica-fore*, and *aeshna-constricta-fore* classes.

In terms of mAP50-95: wing-vein information also performs better in mAP50-95 for the overall class (“all”) and most specific classes such as *aeschna-canadensis-back, aeschna-canadensis-fore, aeshna-constricta-back, aeshna-umbrosa-back, epiaeschna-heros-back*, and *epiaeschna-heros-fore*. On the other hand, wing-outline information exhibits higher mAP50-95 values in *-rhion-aeschna-californica-back* and *-rhion-aeschna-californica-fore*.

In summary, wing-vein information generally exhibits superior performance across most metrics and classes, particularly in recall and mAP metrics, suggesting a more robust and consistent annotation process. According to these validation results, the model indeed utilizes the internal structure of wing veins as informative features, and this structure appears to be uniquely distinguishable for each species. However, for some species, wing-outline information may be more advantageous for species identification.

**Table 2.** Species identification performance with wing vein information (Above) or wing outline information (Below).

| Class | Images | Instances | P | R | mAP50 | mAP50-95 |
| --- | --- | --- | --- | --- | --- | --- |
| all | 41 | 67 | 0.72 | 0.903 | 0.905 | 0.8 |
| rhion-aeschna-californica-back | 3 | 3 | 0.428 | 1.0 | 0.995 | 0.863 |
| rhion-aeschna-californica-fore | 3 | 3 | 0.56 | 1.0 | 0.995 | 0.93 |
| aeschna-canadensis-back | 4 | 4 | 0.803 | 1.0 | 0.995 | 0.885 |
| aeschna-canadensis-fore | 4 | 4 | 0.849 | 1.0 | 0.995 | 0.895 |
| aeshna-constricta-back | 5 | 5 | 0.84 | 1.0 | 0.995 | 0.896 |
| aeshna-constricta-fore | 5 | 5 | 0.692 | 1.0 | 0.938 | 0.866 |
| aeshna-umbrosa-back | 4 | 5 | 0.48 | 0.8 | 0.695 | 0.592 |
| aeshna-umbrosa-fore | 5 | 6 | 0.819 | 0.5 | 0.638 | 0.516 |
| epiaeschna-heros-back | 5 | 5 | 1.0 | 0.964 | 0.995 | 0.869 |
| epiaeschna-heros-fore | 5 | 5 | 0.892 | 0.8 | 0.962 | 0.771 |
| gynacantha-nervosa-back | 11 | 11 | 0.6 | 0.958 | 0.817 | 0.762 |
| gynacantha-nervosa-fore | 11 | 11 | 0.67 | 0.818 | 0.846 | 0.755 |

| Class | Images | Instances | P | R | mAP50 | mAP50-95 |
| --- | --- | --- | --- | --- | --- | --- |
| all | 41 | 67 | 0.618 | 0.8 | 0.812 | 0.752 |
| rhion-aeschna-californica-back | 3 | 3 | 0.778 | 1.0 | 0.995 | 0.913 |
| rhion-aeschna-californica-fore | 3 | 3 | 0.792 | 1.0 | 0.995 | 0.995 |
| aeschna-canadensis-back | 4 | 4 | 0.718 | 1.0 | 0.995 | 0.958 |
| aeschna-canadensis-fore | 4 | 4 | 0.667 | 1.0 | 0.995 | 0.871 |
| aeshna-constricta-back | 5 | 5 | 0.788 | 1.0 | 0.995 | 0.882 |
| aeshna-constricta-fore | 5 | 5 | 0.681 | 1.0 | 0.995 | 0.955 |
| aeshna-umbrosa-back | 4 | 5 | 0.562 | 0.4 | 0.533 | 0.526 |
| aeshna-umbrosa-fore | 5 | 6 | 0.322 | 0.167 | 0.412 | 0.375 |
| epiaeschna-heros-back | 5 | 5 | 0.468 | 0.712 | 0.686 | 0.594 |
| epiaeschna-heros-fore | 5 | 5 | 0.385 | 0.6 | 0.662 | 0.603 |
| gynacantha-nervosa-back | 11 | 11 | 0.645 | 0.818 | 0.706 | 0.673 |
| gynacantha-nervosa-fore | 11 | 11 | 0.614 | 0.909 | 0.779 | 0.68 |

### 3.3 Classification Within Groups After Grouping by Size

**Figure 8.**
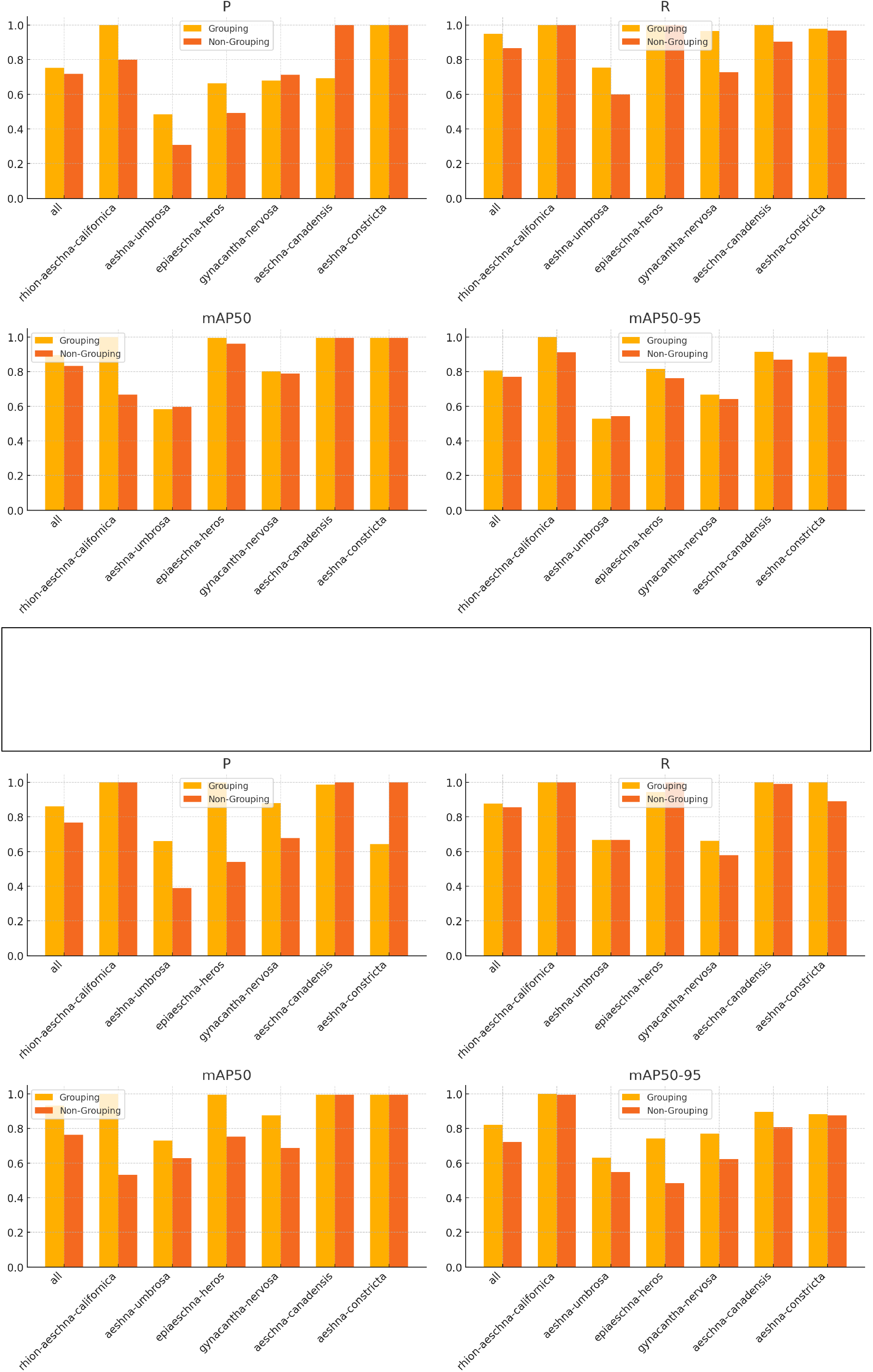
Performance comparison between the non-grouping method vs. classification within groups after grouping by size. The upper graphs represent the performance for the back wings, while the lower graphs show the performance for the fore wings.

The impact of grouping by size on the classification performance of dragonfly species was evaluated. The comparison was conducted between two approaches: a non-grouping method, where all species were classified without considering size, and a method involving classification within groups after grouping by size.

For back wings, the classification performance generally improved when species were first grouped by size before classification. For instance, the Precision (P) for the “all” class improved from 0.719 in the non-grouping method to 0.753 in the grouping method. Similarly, the Recall (R) increased from 0.867 to 0.949. These improvements are reflected across most individual species, particularly for epiaeschna-heros and aeschna-canadensis, which showed significant increases in mAP50 from 0.962 to 0.995 and 0.995 to 0.995, respectively.

Despite these general trends, some species exhibited minor decreases in certain metrics. For example, the mAP50-95 for rhion-aeschna-californica slightly dropped from 0.913 in the non-grouping method to 0.806 in the grouping method. However, the overall improvement across metrics suggests that the grouping method enhances the model’s ability to accurately classify species based on their back wings.

For fore wings, similar to the back wings, the grouping method generally resulted in better classification metrics. The Precision (P) for the “all” class increased from 0.768 to 0.861, while Recall (R) saw a moderate improvement from 0.855 to 0.878. The mAP50 for the “all” class also improved significantly, from 0.765 in the non-grouping method to 0.932 in the grouping method.

For individual species, such as aeschna-constricta, the metrics remained high across both methods, but with a notable improvement in mAP50-95 from 0.875 in the non-grouping approach to 0.883 in the grouping method. However, for gynacantha-nervosa, the improvement was more moderate, with mAP50-95 only slightly increasing from 0.623 to 0.77.

**Table 3.** Performance comparison between the non-grouping method vs. classification within groups after grouping by size. The upper table represent the performance for the back wings, while the lower table show the performance for the fore wings.

| **Class (fore wing; grouping)** | **Images** | **Instances** | **P** | **R** | **mAP50** | **mAP50-95** | **Group** |
| --- | --- | --- | --- | --- | --- | --- | --- |
| all | 33 | 34 | 0.861 | 0.878 | 0.932 | 0.820 | *grouping* |
| rhion-aeschna-californica | 3 | 3 | 1.0 | 1.0 | 1.0 | 1.0 | *grouping* |
| aeshna-umbrosa | 5 | 6 | 0.659 | 0.667 | 0.732 | 0.631 | *grouping* |
| epiaeschna-heros | 5 | 5 | 1.0 | 0.944 | 0.995 | 0.743 | *grouping* |
| gynacantha-nervosa | 11 | 11 | 0.879 | 0.661 | 0.875 | 0.77 | *grouping* |
| aeschna-canadensis | 4 | 4 | 0.988 | 1.0 | 0.995 | 0.895 | *grouping* |
| aeshna-constricta | 5 | 5 | 0.644 | 1.0 | 0.995 | 0.883 | *grouping* |
| all | 33 | 34 | 0.768 | 0.855 | 0.765 | 0.722 | *non-grouping* |
| rhion-aeschna-californica | 3 | 3 | 1.0 | 1.0 | 0.532 | 0.995 | *non-grouping* |
| aeshna-umbrosa | 4 | 4 | 0.39 | 0.667 | 0.629 | 0.548 | *non-grouping* |
| epiaeschna-heros | 5 | 5 | 0.541 | 1.0 | 0.753 | 0.484 | *non-grouping* |
| gynacantha-nervosa | 5 | 6 | 0.679 | 0.58 | 0.687 | 0.623 | *non-grouping* |
| aeschna-canadensis | 5 | 5 | 1.0 | 0.991 | 0.995 | 0.808 | *non-grouping* |
| aeshna-constricta | 11 | 11 | 1.0 | 0.89 | 0.995 | 0.875 | *non-grouping* |

| **Class (fore wing; grouping)** | **Images** | **Instances** | **P** | **R** | **mAP50** | **mAP50-95** | **Group** |
| --- | --- | --- | --- | --- | --- | --- | --- |
| all | 33 | 34 | 0.753 | 0.949 | 0.895 | 0.806 | *grouping* |
| rhion-aeschna-californica | 3 | 3 | 1.0 | 1.0 | 1.0 | 1.0 | *grouping* |
| aeshna-umbrosa | 4 | 5 | 0.484 | 0.755 | 0.584 | 0.528 | *grouping* |
| epiaeschna-heros | 5 | 5 | 0.664 | 1.0 | 0.995 | 0.816 | *grouping* |
| gynacantha-nervosa | 11 | 11 | 0.68 | 0.966 | 0.803 | 0.667 | *grouping* |
| aeschna-canadensis | 4 | 4 | 0.693 | 1.0 | 0.995 | 0.914 | *grouping* |
| aeshna-constricta | 5 | 5 | 1.0 | 0.978 | 0.995 | 0.911 | *grouping* |
| all | 32 | 33 | 0.719 | 0.867 | 0.834 | 0.770 | *non-grouping* |
| rhion-aeschna-californica | 3 | 3 | 0.798 | 1.0 | 0.667 | 0.913 | *non-grouping* |
| aeshna-umbrosa | 4 | 4 | 0.309 | 0.6 | 0.595 | 0.542 | *non-grouping* |
| epiaeschna-heros | 5 | 5 | 0.492 | 1.0 | 0.962 | 0.763 | *non-grouping* |
| gynacantha-nervosa | 4 | 5 | 0.713 | 0.727 | 0.789 | 0.642 | *non-grouping* |
| aeschna-canadensis | 5 | 5 | 1.0 | 0.903 | 0.995 | 0.871 | *non-grouping* |
| aeshna-constricta | 11 | 11 | 1.0 | 0.969 | 0.995 | 0.887 | *non-grouping* |

## 4. Conclusion

This study investigates the digitization of insect collections and the application of deep learning models to improve this process. During a one-hour filming session, 141 images of dragonfly specimens from the Cornell University Insect Collection were captured and preprocessed using five distinct methods: (1) adding box annotations around the wings, (2) adding polygon annotations to outline the forewings and hindwings, (3) removing vein system images, (4) retaining only the wing outline images, and (5) grouping by automatically measured wing size and classifying species within those groups. By comparing YOLOv8 models trained on datasets with these different preprocessing methods, the study revealed three key findings: (1) datasets with bounding box annotations result in shorter preprocessing times and superior model performance compared to polygon annotations; (2) although models trained with polygon annotations may have lower accuracy, polygon annotations provide more detailed information on wing length and phenotypic traits; and (3) the wing vein system, rather than the wing outline, is the critical factor in classification accuracy.

